# Context and Attention Shape Electrophysiological Correlates of Speech-to-Language Transformation

**DOI:** 10.1101/2023.09.24.559177

**Authors:** Andrew J. Anderson, Christopher Davis, Edmund C. Lalor

**Affiliations:** Department of Neurology. Medical College of Wisconsin. 9200 W. Wisconsin Ave. Milwaukee, WI 53226 USA.; Department of Neuroscience and Del Monte Institute for Neuroscience, University of Rochester, 201 Robert B. Goergen Hall, P.O. Box 270168, Rochester, NY 14627, USA; Western Sydney University, The MARCS Institute for Brain, Behaviour and Development, Westmead Innovation Quarter, Building U, Level 4, 160 Hawkesbury Road, Westmead NSW 2145. Australia; Department of Biomedical Engineering and Center for Visual Science, University of Rochester, 201 Robert B. Goergen Hall, P.O. Box 270168, Rochester, NY 14627, USA.

## Abstract

To transform speech into words, the human brain must accommodate variability across utterances in intonation, speech rate, volume, accents and so on. A promising approach to explaining this process has been to model electroencephalogram (EEG) recordings of brain responses to speech. Contemporary models typically invoke speech categories (e.g. phonemes) as an intermediary representational stage between sounds and words. However, such categorical models are typically hand-crafted and therefore incomplete because they cannot speak to the neural computations that putatively underpin categorization. By providing end-to-end accounts of speech-to-language transformation, new deep-learning systems could enable more complete brain models. We here model EEG recordings of audiobook comprehension with the deep-learning system Whisper. We find that (1) Whisper provides an accurate, self-contained EEG model of speech-to-language transformation; (2) EEG modeling is more accurate when including prior speech context, which pure categorical models do not support; (3) EEG signatures of speech-to-language transformation depend on listener-attention.

## Introduction

The apparent ease with which the human brain transforms speech sounds into words belies the complexity of the task. This complexity is due in large part to speech variability - each time a word is spoken, the sound is different. Speech variability is most striking in extreme cases such as when people have unfamiliar accents, shout, whisper or sing, but is always present to some degree, even when the same person repeats the same phrase (Smith et al. 1995). How brains transform such variable speech sounds into language is a key unresolved question in cognitive neuroscience. By enabling temporally precise estimates of brain activity, electrophysiological measures such as EEG have provided evidence that the brain transforms natural continuous speech into words as a cascading process, with abstract categorical speech units such as phonemes serving as intermediary pre-lexical representations (di Liberto et al. 2015, and very recently Gillest et al. 2023). This evidence has however been challenged (Daube et al. 2019) due to its reliance on so-called oracle models (Kriegeskorte 2018) of pre-lexical representation that categorize speech sounds into time-series of hand-crafted sub-lexical units. Such approaches are limited by: (1) Providing no explicit computational model of the brain mechanism that putatively categorizes speech units (which means they cannot capture associated brain activity); (2) Modelling information that was not available to participants’ brains during listening (i.e. speech categories); and (3) Requiring manual intervention to code speech units and verify time-alignment. The current study helps to relieve these limitations by modeling EEG recordings of natural speech, with high accuracy, using a state-of-the-art deep-learning speech-to-language model (Whisper, Radford et al. 2022). Critically Whisper computes a graded transformation of speech into language without experimenter intervention.

A second contribution of the study is to question a common modelling assumption implicit in previous work that has either supported (di Liberto et al. 2015) or challenged (Daube et al. 2019) evidence that electrophysiological speech responses reflect abstract speech representation as opposed to spectral features. The implicit assumption is that low-level electrophysiological speech responses are largely context-invariant, in that they echo concurrent speech features with delays of up to 300ms. When context has been built into neurophysiological models of speech transformation, it has been computed based on within word probabilities of successive phonemes (Brodbeck et al. 2018). In contrast, context sensitivity to previous words and sentences is widely documented in language-related responses such as the N400 (Kutas et al. 1980, 2011) which is prominent in centroparietal electrodes ∼400ms post word onset. Despite evidence on either side (McClelland and Elman, 1986; Norris et al., 2000; Davis and Johnsrude, 2007; Travis et al., 2013), one prevailing view is that high-level linguistic context predictively shapes pre-lexical representation (Kuperberg and Jaeger, 2016), which has received some electrophysiological support in natural speech comprehension (e.g. Broderick et al. 2019). We here present evidence that EEG responses in bilateral temporal electrodes that are typically linked to low-level speech processing are sensitive to longer range speech contexts, and in particular that incoming speech sounds may be refined and integrated with past speech heard up to 10s back. Encoding prior context could help the brain leverage language to anticipate speech sounds and disambiguate periods of noisy speech (e.g. given “President Joe <NOISE>” a good guess is that the noise obscures “Biden”). Such contextual information is not present in purely categorical speech models.

To support the above claims, we modeled EEG recordings taken as participants listened to audiobooks based on the activation patterns of speech-to-language deep-learning system (Whisper). Whisper exploits speech context to transform spectral speech features into a time-aligned series of linguistic vectors. A noteworthy characteristic of the contextualized transformation is that it is graded across a series of feedforward processing layers, which presented us with the opportunity to examine EEG sensitivity to different stages of speech-to-language transformation. Specifically, each successive Whisper layer uses so-called self-attention computations to contextualize each input time-frame (either input spectral features or the previous layer output) as a weighted composite of itself and previous time-frames (up to 30s back). We find that Whisper’s more linguistic contextualized layers predict EEG with high accuracy, above and beyond early layers and other competing models of speech acoustics and language. We further consolidate this evidence with a follow up analysis of a second Cocktail Party selective-attention dataset (O’Sullivan et al. 2014). Here participants heard two concurrent speakers, and selectively paid attention to one – which we hypothesized, would selectively reflect the speech-to-language transformation performed by Whisper (as we later find).

EEG recordings of audiobook comprehension were analyzed. The audiobook waveform was processed through a pre-trained deep-learning speech-to-language model (Whisper-base). A sliding window approach was applied to feed the model up to 30s of prior speech audio waveform, which was then re-represented as an 80 channel Log-Mel Spectrogram. The Spectrogram is then fed-forward through successive layers of a Transformer Encoder artificial neural network via an initial convolutional layer. This entire process can be considered to implement a graded transformation of input speech to a contextualized linguistic representation. At each transformer layer, input time-frames are contextualized via a self-attention computation that re-represents input frames according to a weighted average of themselves and preceding frames. The bottom row illustrates a summary of self-attention weightings computed at each layer for the first 3s of the audiobook stimulus. Attention weights relating each time frame to each previous time-frame are illustrated as the colored shading on each matrix row. Specifically, points along the diagonal correspond to attention weights at any timepoint t (from 0 to 3s) unrelated to preceding context. Meanwhile, points to the left of the diagonal correspond to the attention weights applied to preceding frames to re-represent the current time frame. The wealth of color to the left of the diagonal in layers 3, 5, and 6 demonstrates the importance of prior context in Whisper’s operation. The self-attention computation is illustrated in detail in **Supp. Fig. 1**, and self-attention weight maps computed in the eight attention heads that were summed to generate the visualization above are in **Supp Fig. 2/3**. Whisper layer outputs were used to predict co-registered EEG data in a cross-validated multiple regression framework (this is illustrated above for only the final layer output). To reduce computational burden, Whisper vectors were reduced to 10 dimensions by projection onto pre-derived PCA axes (computed from different audiobook data, see also **Supp Fig. 4**), and both EEG and model data were resampled at 32Hz.

## Results: Overview

The overarching aims of the forthcoming analyses were twofold: (1) To establish how EEG recordings of speech comprehension reflect Whisper’s graded linguistic transformation of speech, both in natural listening conditions and when speech is paid attention to or not. Here, we anticipated that unattended speech would be processed at a more superficial level, and thus EEG correlates of linguistic transformation would dwindle. (2) To estimate how elements of context, speech and language encoded within Whisper contributed to EEG prediction.

We first reanalyzed a dataset of publicly available Electroencephalographic (EEG) recordings (https://doi.org/10.5061/dryad.070jc) taken from 19 subjects as they listened to ∼1hour of an audiobook (The Old Man and the Sea, Hemingway, 1952). We hypothesized that the internal representations of Whisper – which reflect a graded transformation of spectral speech input into language – would more accurately predict EEG responses than spectral features of the speech, or derivatives thereof. This was because Whisper, like the human brain, is adapted to transform speech to language. Because it is well established from N400 studies that EEG is sensitive to language (Kutas et al. 1980, 2011) and in particular word expectation (Frank et al. 2015, Heilbron et al. 2022), we ran a battery of control analyses to gain confidence that Whisper was not re-explaining established signatures of language processing. These control tests were primarily based on estimates of lexical (word) surprisal (how unexpected a word is based on prior context), though we later reference findings to the internal states of the language model used to generate next word expectations (GPT-2, Radford et al. 2019).

## Natural Speech EEG Recordings Strongly Reflect the Linguistic Transformation of Acoustic Speech

To establish how accurately different Whisper layers predicted EEG and determine if they complemented acoustic and lexical models of speech, we ran a series of cross-validated multiple regression analyses. In each analysis, a model to EEG mapping was fit to each individual’s data. To estimate the complementary predictive value of different models, EEG prediction accuracies derived from model combinations were contrasted with those from constituent models. Model combinations are referred to as Unions in forthcoming text and/or completely specified as [Whisper LX Control], where LX corresponds to Layer number X and the square brackets indicate concatenation of Whisper features with coincident control model features. Control models were: (a) The audiobook speech envelope concatenated with its first half wave rectified derivative (abbreviated as Env&Dv). (b) An 80 channel Log-Mel Spectrogram corresponding to Whisper’s input and computed by Whisper’s preprocessing module. (c) Lexical surprisal – a measure of how unexpected each word is. Surprisal values were computed using GPT-2 to anticipate the identity of each forthcoming word based on up to 1024 prior words, which is represented as a long vector of probability values linked to each word in GPT-2’s dictionary. Lexical surprisal values for individual words were computed as the negative log probability of the actual word_n+1_. A time-series representation was then constructed by aligning lexical surprisal “spikes” to word onset times (relative to the EEG time-line) and setting all other time-series values to zero.

To first establish whether Whisper complemented each individual control model, we evaluated whether pairwise [Whisper LX Control] scalp-average EEG predictions were more accurate than predictions derived from the Control alone. This evaluation used signed-ranks tests (one-tail, 19 subjects), with p-values corrected across layers for multiple comparisons using False Discovery Rate (FDR). There were two principal findings (both illustrated in Figure 2 **Top Row**): (1) Whisper vectors from L1-L6 strongly complemented all Controls in prediction. For example, [Whisper L6 Env&Dv] had an accuracy of Mean±SEM r=0.056±0.004, which doubled the variance predicted by Env&Dv (r=0.041±0.003) - where predicted variance was computed as r^2^. (2) EEG prediction accuracies became successively stronger as a function of layer depth. The Mean±SEM Spearman correlation coefficient between prediction accuracy and layer depth (0 to 7) across participants was 0.77±0.07. The set of correlation coefficients were significantly greater than 0 (Signed-rank Z=3.7924, p=1.5e-4, n=19, 2-tail). This provided evidence that deeper and more linguistic Whisper layers were the strongest EEG predictors.

**Figure 1.**
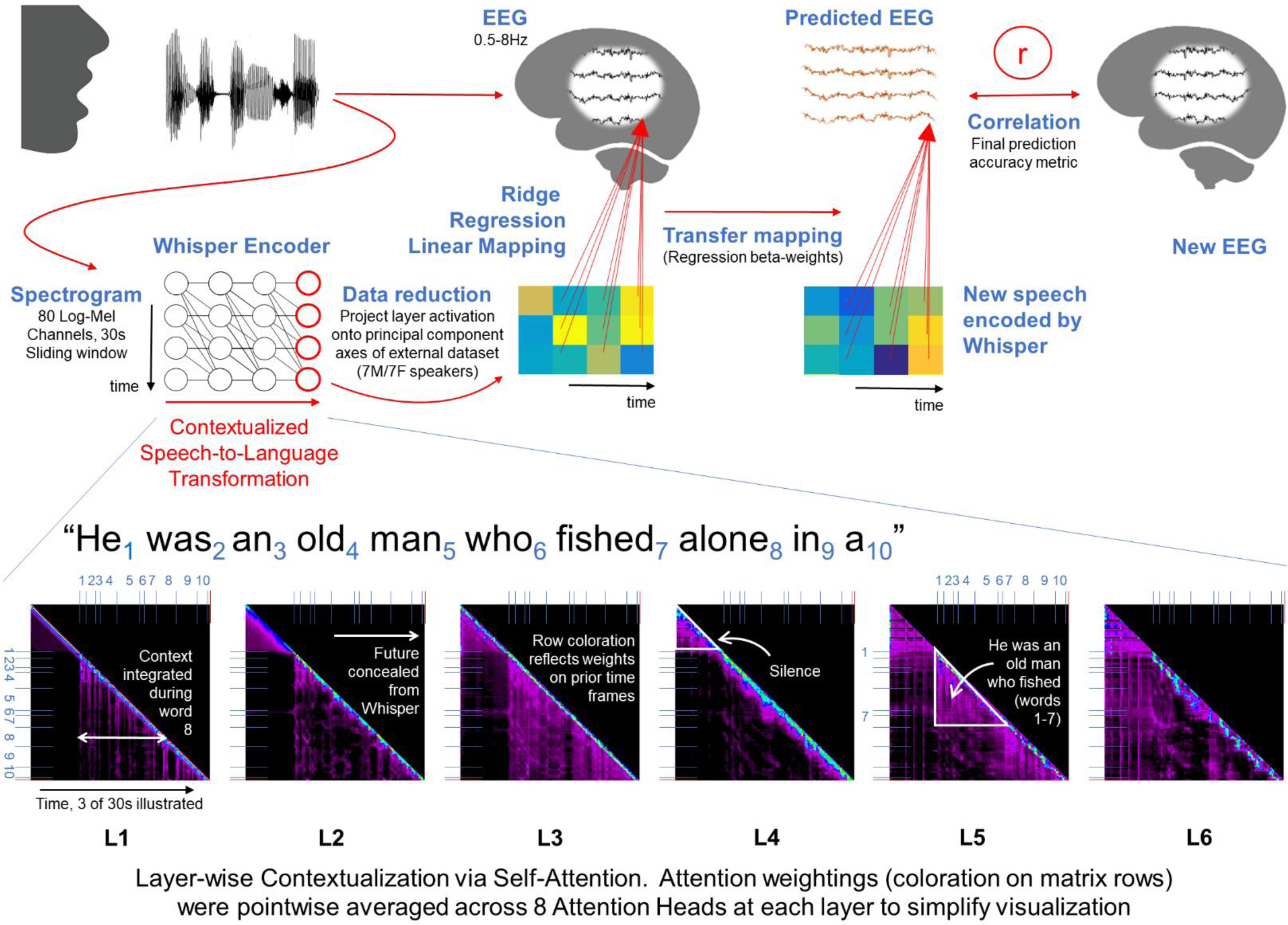
Predicting Natural Speech EEG Recordings with a Contextualized Speech-to-Language Model

**Figure 2.**
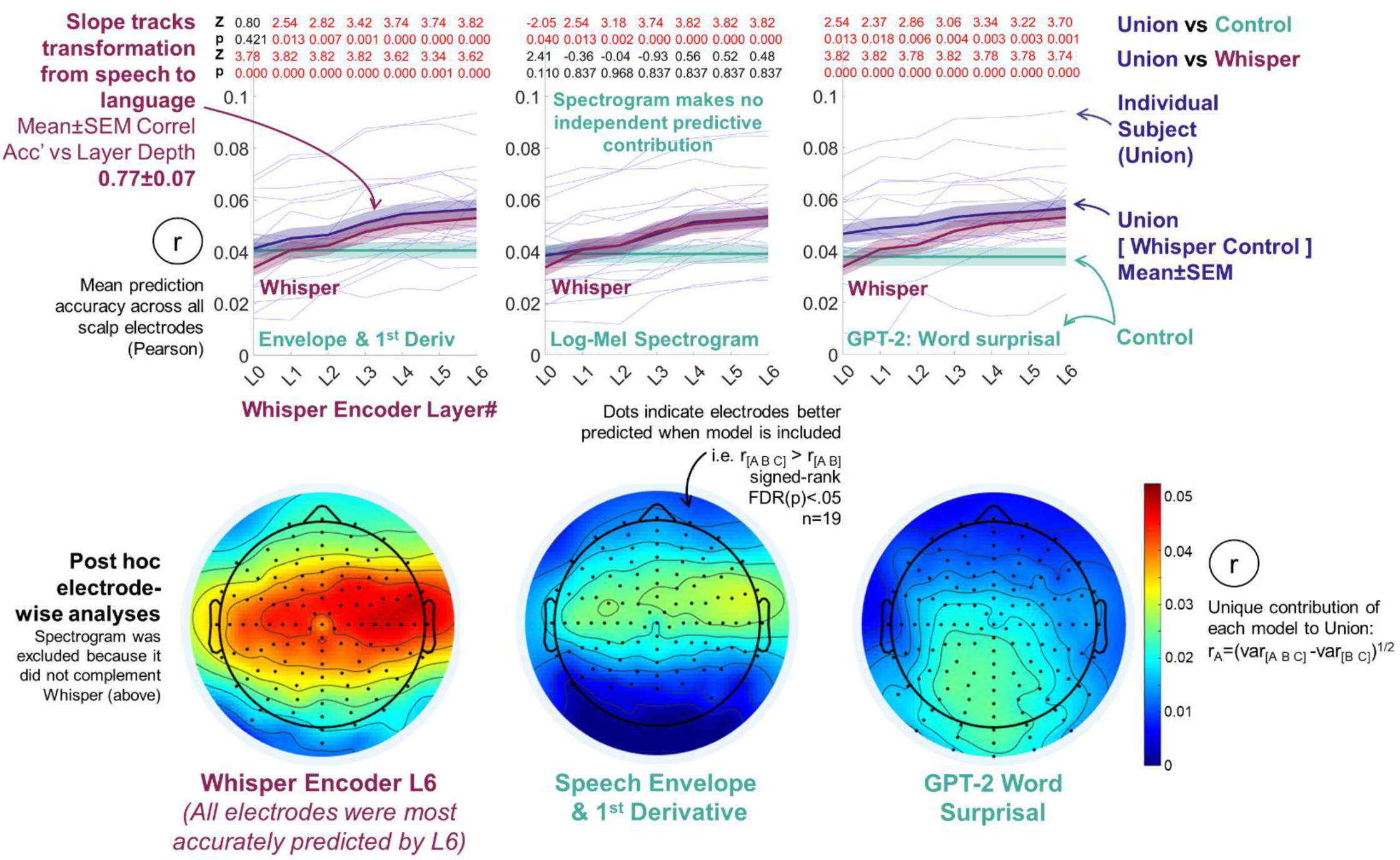
EEG is more accurately predicted by a contextualized speech-to-language model than standard acoustic or lexical surprisal representations, with accuracy increasing with model layer depth. Top row plots: The speech-to-language model (Whisper) complemented Speech Envelope-based measures, a Log-Mel Spectrogram (Whisper’s input) and lexical surprisal in predicting EEG data. Models were considered to be complementary if scalp-average EEG prediction derived from concatenating model pairs (Union=[Whisper, Surprisal]) were more accurate than constituent models (signed-ranks tests, Z and FDR corrected p-values are displayed at the top of each plot). Whisper Layers 1 to 6 complemented every competing predictor. Lexical surprisal and the envelope-based predictors, but not the spectrogram also made independent contributions to prediction. Corresponding test statistics (Z, FDR(p)) are in the first and second row of each plot). Bottom row scalp maps: Post hoc electrode-wise analyses mapped scalp-regions that were sensitive to the speech-to-language model. Each electrode was predicted using the Union of Whisper Layer 6, the Envelope-based measures and Lexical Surprisal (because all three had made independent predictive contributions in the primary scalp-average analyses, unlike the spectrogram which was excluded). Whisper’s independent contribution was estimated by partitioning the variance predicted by the three-model union (including Whisper) minus the variance predicted by a joint model excluding Whisper (Envelope and Surprisal). The square root of the resultant variance partition was taken to provide a pseudo-correlation estimate on the same scale as other results, as is a common procedure (de Heer et al. 2017, Vaidya et al. 2022). Electrodes that are typically associated with low-level acoustic processing were especially sensitive to Whisper. This is visible as the red band that straddles central scalp from ear to ear in the leftmost scalp map.

To identify which Control models complemented Whisper, we evaluated whether Union prediction accuracies were greater than constituent Whisper layers (Signed ranks tests, one-tail, 19 subjects). This revealed that both Env&Dv and Lexical Surprisal, but not the Spectrogram (Whisper’s input) uniquely predicted elements of the EEG signal. In sum, these results provide evidence that EEG strongly reflects the transformation of speech to language encoded by Whisper.

To next establish which electrodes reflected speech-to-language transformations, we ran a series of post hoc electrode-wise prediction analyses that combined Whisper’s final most linguistic layer (L6 – which as it turned out also generated the most accurate predictions of each electrode) with both Env&Dv and Lexical Surprisal. The Log-Mel Spectrogram was excluded from this and all further analyses because it had made no independent predictive contribution in our initial analyses (Figure 2 **Top Row**). To estimate whether Whisper uniquely contributed to predicting an individual electrode, signed ranks tests (19 subjects, one-tail) were used to evaluate whether electrode predictions derived from three model Union: [Whisper L6 Env&Dv Lexical Surprisal] were more accurate than when Whisper was excluded (i.e., [Env&Dv Lexical Surprisal]). P-values across the 128 channels were FDR corrected for multiple comparisons. Analogous tests were deployed to reveal each electrode’s unique sensitivity to Env&Dv (e.g. Union vs [Whisper Lexical Surprisal]) as well as Lexical Surprisal (Union vs [Whisper Env&Dv]). To estimate the relative contribution of each model to predicting each electrode we deployed a predicted-variance partitioning approach (de Heer et al. 2017, Vaidya et al. 2022). For instance, Whisper’s independent predictive contribution was estimated as the variance predicted by the Union minus the variance predicted when Whisper was excluded, i.e. r_2[Whisper Env&Dv Lexical Surprisal]_ - r_2[Env&Dv Lexical Surprisal]_. The square root of the resultant variance partition was taken to provide a pseudo-correlation estimate, as is a common practice. In the instance of negative variance partitions (as can arise from overfitting in regression, despite regularization), pseudo correlation estimates were zeroed. Scalp maps of pseudo correlation variance partition estimates overlaid with the outcomes of the above signed ranks tests are illustrated in Figure 2 **Bottom Row**.

The electrode-wise analyses revealed two principal findings: (1) Linguistic Whisper L6 dominated prediction in bilateral temporal scalp electrodes that are traditionally associated with low-level speech processing (and also captured here in part by Env&Dv). (2) EEG signatures of Lexical Surprisal were distinct and observed in the centroparietal electrodes that are traditionally associated with the N400. In sum, these results link the new EEG correlates of speech-to-language transformation to bilateral temporal scalp regions which are typically associated with processing acoustic speech or speech units.

## EEG Correlates of Speech-to-Language Transformation Are Attention-Dependent in Cocktail-Party Environments

Given that Whisper accurately predicts scalp EEG in traditional speech processing regions, it was possible that Whisper’s predictive advantage was gained by providing a high-quality brain-like filter of concurrent surface-level spectral features. We reasoned that one compelling test of this would be to modulate the depth of linguistic processing on the listener’s side, and test whether the EEG data’s sensitivity to Whisper was likewise modulated.

One effective way of modulating a listener’s engagement to speech is via selective attention. In so-called cocktail-party scenarios it is widely appreciated that listeners can home in on a single speaker whilst ignoring others (Mesgarani and Chang 2012, Ding and Simon 2012, O’Sullivan et al. 2014). Indeed, the electrophysiological bases of selective attention in multi-talker environments have been thoroughly investigated and there is general agreement that unattended speech is processed at a more superficial level than attended speech (Mesgarani and Chang 2012, Ding and Simon 2012, O’Sullivan et al. 2014, Broderick et al. 2018). For instance, listeners have low ability to accurately report on the unattended speech content, and N400-like lexical expectation responses either dwindle or disappear for unattended speech whilst traces of low-level acoustic speech processing remain (Broderick et al. 2018), albeit with a reduced magnitude. We therefore hypothesized that the new indices of speech-to-language transformation observed in Figure 2 would be present for an attended speaker but would dwindle or disappear for unattended speech. In particular, we hypothesized that the predictive advantage associated with Whisper layer-depth (the slope across layers in Figure 2) would flatten out for unattended speech.

To test the above hypothesis, we reanalyzed a publicly available Cocktail Party EEG dataset (https://doi.org/10.5061/dryad.070jc) where EEG was recorded as listeners were played 30mins of two audiobooks presented simultaneously. The two audiobooks – “20,000 Leagues under the Sea” and “Journey to the Center of the Earth” were narrated by different male speakers, and were presented via headphones, one to the left ear and the other the right. Participants were tasked with selectively attending to one audiobook (in one ear) across the entire experiment. We analyzed 15 participants who paid attention to “20,000 Leagues…” and 12 who paid attention to “Journey…”. Audiobook presentation was split in thirty runs of 1 min each that were separated by brief breaks to mitigate participant fatigue. 6 participants in the online data repository with incomplete EEG data sets were excluded from forthcoming analyses. This enabled the analyses to be standardized to be have exactly the same parameters for each person to support precise comparisons.

To establish how speech-to-language signatures varied with listener attention, we repeated the battery of analyses presented in Figure 2 whilst modelling either the attended or unattended speech stream. Besides predicting the two speech streams, the only other difference to our first analyses was that we discontinued the using the Log-Mel spectrogram because it had previously afforded no predictive benefit (Figure 2).

For attended speech, as is illustrated in Figure 3 **Upper Left**, the entire pattern of scalp-average EEG prediction accuracies for the different pairwise model combinations broadly corroborated the earlier finding that the deeper more linguistic Whisper layers were more accurate EEG predictors (Figure 2). Specifically, signed ranks comparisons of scalp-average prediction accuracies revealed that combining Whisper L4-6 with Env&Dv elevated prediction accuracies above Env&Dv, and all Whisper layers improved significantly over Lexical Surprisal (See Figure 3 for test statistics). For example, [Whisper L6 Env&Dv] had a Mean±SEM accuracy of r=0.055±0.003 comparative to Env&Dv (r=0.041±0.003) where Env&Dv was the most accurate Control. Also echoing Figure 2, prediction accuracies increased with layer depth – and speech-to-language transformation. The Mean±SEM Spearman correlation between prediction accuracy and layer depth was 0.78±0.06, which was significantly greater than zero (Signed-rank Z=4.4, p=1.2e-5, n=27, 2-tail).

**Figure 3.**
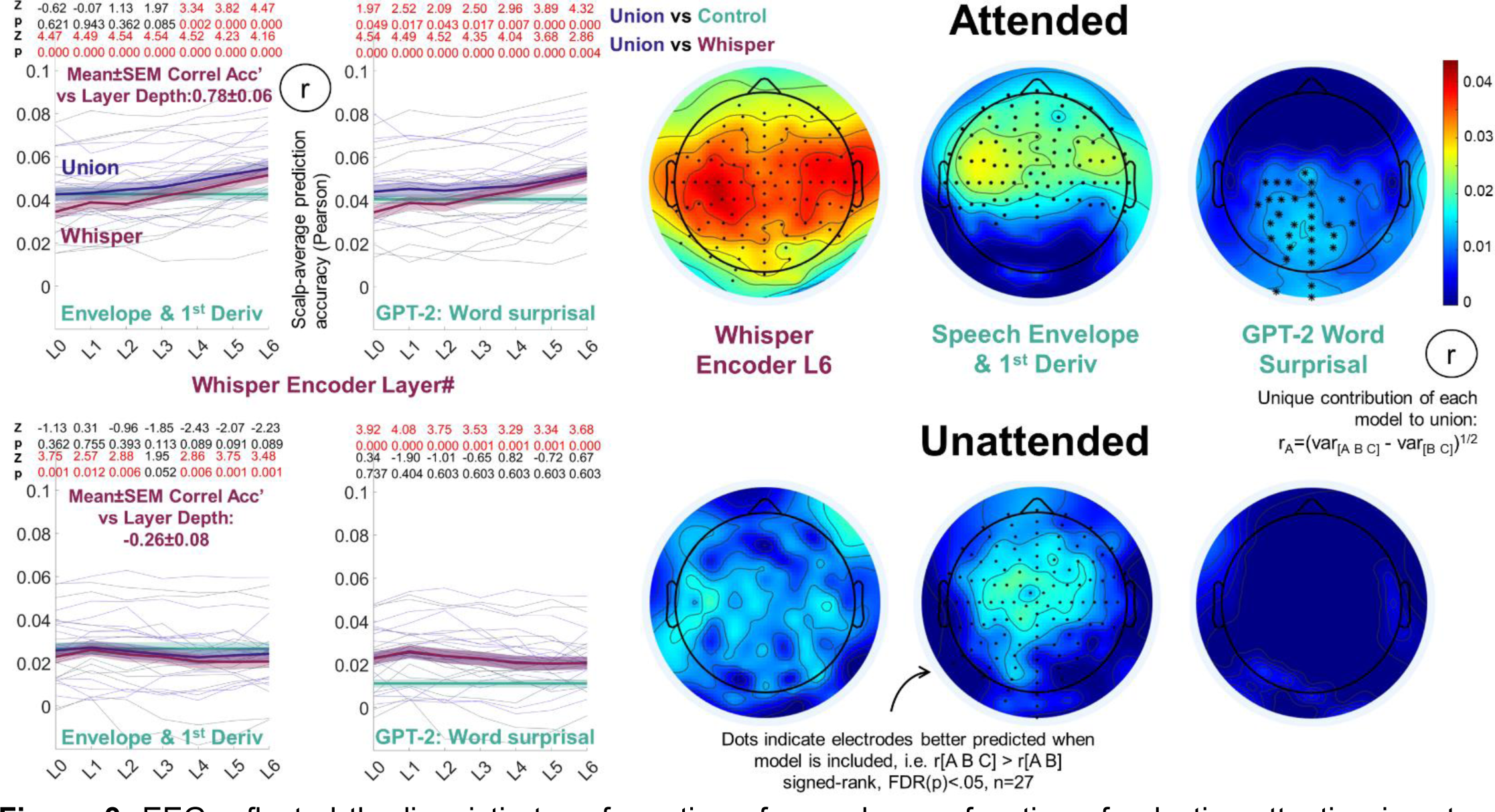
EEG reflected the linguistic transformation of speech as a function of selective-attention in a two-speaker cocktail party scenario. **Top Row:** EEG Prediction accuracies derived from attended speech models resembled the single audiobook data (Figure 2). **Top Left Plots:** The speech-to-language model (Whisper) complemented speech envelope-based measures and lexical surprisal in scalp-average EEG prediction. Z and FDR corrected p-values at the top of each plot correspond to signed ranks tests comparing Union EEG prediction accuracies to constituent models. Mirroring Figure 2, Whisper’s prediction accuracy increased with layer depth and latter layers complemented both the envelope measures and lexical surprisal. **Top Right Scalp Maps:** Electrode-wise analyses and variance partitioning (see Figure 2 caption for details) revealed that Whisper L6 dominated prediction in bilateral temporal scalp regions. Lexical surprisal was reflected in centroparietal electrodes. **Bottom Row:** EEG Prediction accuracies derived from unattended speech models revealed no linguistic contribution. **Bottom Left Plots:** Based on scalp-average measures, Whisper added no predictive value to the envelope-based measures, and rather than increasing, prediction accuracy was weaker in later layers (see signed ranks Z and FDR corrected p-values at the top of each plot). Whisper did however improve on lexical surprisal prediction accuracies (all layers), which is presumably because it still encodes a residual of unattended speech acoustics. **Bottom Right Scalp Maps:** Consistent with the brain processing only superficial acoustic features of unattended speech electrode-wise analyses and predicted variance partitioning echoed that envelope-based measures alone drove prediction over central scalp electrodes.

Follow up electrode-wise analyses that partitioned how Whisper L6, Env&Dv and Lexical Surprisal each contributed to predicting attended speech revealed scalp maps (Figure 3 **Upper Right)** that again echoed Figure 2. Whisper L6 dominated responses in bilateral temporal electrodes, Env&Dv also contributed to predicting electrodes in those same scalp locations, and Lexical Surprisal made independent contributions to predicting centroparietal electrodes. The most salient difference to Figure 2 was that the early Whisper layers (pre-L4) failed to improve over Env&Dv predictions. This could reflect the more challenging listening conditions, or contamination of the EEG signal with traces of acoustic processing of unattended speech.

Differently, and in line with the hypotheses that EEG correlates of speech-to-language transformation would dwindle for unattended speech, no Whisper model of unattended speech improved on the scalp-average prediction accuracies made by Env&Dv (Figure 3 **Lower Left**). Whisper did however complement Lexical Surprisal and this effect was observed for all Whisper layers. Based on our electrode-wise analyses, we are confident that this effect stems from traces of speech acoustics that are residual in Whisper (which is derived directly from spectral speech features) but absent from Lexical Surprisal time-series (which is divorced from speech acoustics aside from tracking word onset times). Specifically, follow up electrode-wise analyses comparing [Whisper L6 Env&Dv Lexical Surprisal] to two-model subsets (e.g. [Whisper L6 Lexical Surprisal]), revealed that Env&Dv was the only representation to uniquely contribute to EEG prediction, and this effect was observed in frontocentral electrodes (Figure 3 **Bottom Right**). Thus Env&Dv in isolation provided the most parsimonious EEG model of unattended speech.

Our second selective-attention hypothesis was that when unattended speech was modelled, Whisper layer depth would have little influence on prediction accuracy (the slope across layers in Figure 2 would flatten). As it turned out, the Mean±SEM Spearman correlation between Whisper prediction accuracy and layer depth across participants was negative: -0.26±0.08, and significantly beneath zero when tested across participants (Signed-rank Z=-2.6, p=0.01, n=27, 2-tail). Thus, rather than increasing with layer depth and the transformation from speech-to-language (as for attended speech) EEG prediction accuracy decreased. To provide extra support for this claim, **Supplementary Table 1** presents a linear mixed model analysis of the entire Selective-Attention data set of scalp-average prediction accuracies displayed in Figure 3.

In sum, the analyses of cocktail party selective-attention EEG data provide evidence that when listeners disengage from speech – and traditional N400 signatures of lexical processing diminish, the predictive advantage afforded by Whisper in bilateral temporal electrodes also diminishes. This is consistent with the claim that Whisper recovers an electrophysiological signature of speech-to-language transformation (when speech is attended to at least) rather than providing a brain-like filter of concurrent spectral features (which are still present when speech is not attended).

## Interpreting EEG Correlates of Speech-to-Language Transformation

Having consolidated evidence that EEG indexes linguistic transformations of speech, we finally sought to probe the nature of the new EEG signal predicted and in particular estimate the extent to which it reflected lexical processing and contextualization. We focused this investigation on the single audiobook EEG dataset (Figure 2), because EEG recordings were longer and not contaminated with dual speech streams unlike the Cocktail Party data set.

## Interpretation: EEG Correlates of Whisper Partially Reflect Predictive Lexical Processing

Because Whisper’s last and putatively most linguistic layer was universally the most accurate EEG predictor, we ran further tests to explore how strongly the EEG correlates reflected lexical processing. To this end we first referenced Whisper’s EEG predictions to the internal states of a pure language model (GPT-2). To recap, GPT-2 also underpinned the earlier lexical surprisal analysis, and was chosen due to its excellent universal performance in predicting brain data (Sun et al. 2020, Schrimpf et al. 2021, Caucheteux et al. 2022, Goldstein et al. 2022, Heilbron et al. 2022). The current analysis differs in its basis on GPT-2’s internal states – or layer activations – which are generally thought to encode a mixture of lexical semantics and syntax that help GPT-2 to predict upcoming words. To contrast Lexical Surprisal (Figure 2 **and 3**) reflects the inaccuracy of GPT-2’s next-word-predictions (which is the error signal used to optimize GPT-2).

To establish commonalities and complementarities of GPT-2 with different Whisper layers, we predicted EEG with pairwise combinations of GPT-2 L16 and each Whisper layer and then examined how scalp-average prediction accuracy compared to the two isolated constituent models. We selected L16 based on the outcomes of independent research studies computed on fMRI (Caucheteux et al. 2022), but nonetheless ran post hoc tests with different layers that corroborated the validity of this choice (**Supp Fig 5**).

Results are illustrated in Figure 4 **Left**. Signed ranks comparisons of prediction accuracies revealed that combining GPT-2 with early (L0-4) but not late (L5-L6) Whisper layers boosted accuracy above Whisper alone. Notably, there also were negligible differences in prediction accuracy between Unions of GPT-2 L16 with early and late Whisper layers. For instance the Mean±SEM Spearman Correlation coefficient between [GPT-2 L0-6] scalp-average prediction accuracy and layer depth (L0-6) was 0.14±0.09, which was not significantly greater than zero (Signed rank Z=0.85, p=0.4). This suggests that Whisper L5 and L6 share commonalities in lexical representation with GPT-2 that are lacking from Whisper L0 to L4. Critically, this provides evidence that EEG correlates of Whisper L6 indeed reflect lexical transformations of speech and also support the notion that Whisper encodes a graded speech-to-language transformation (which we had assumed prior to this analysis).

**Figure 4.**
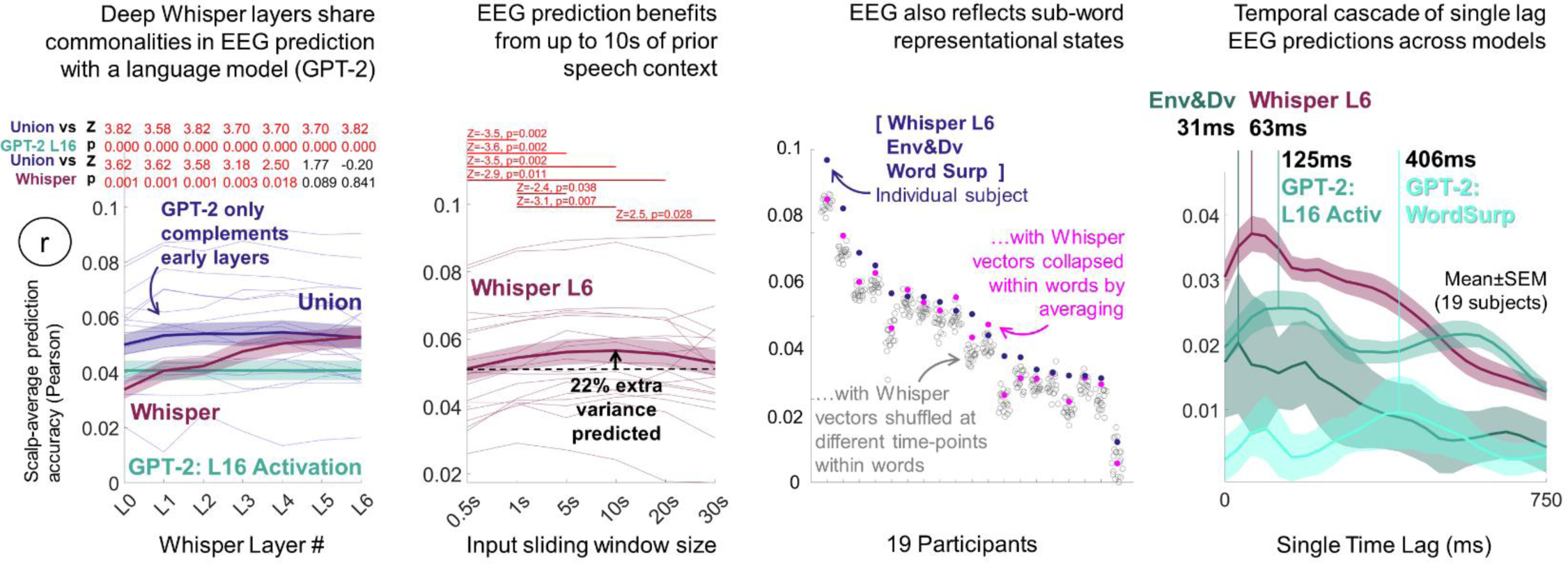
Interpretation: EEG correlates of speech-to-language transformation reflect a blend of contextualized lexical and sub-lexical representation. **Left:** To examine the linguistic nature of Whisper’s EEG predictions we referenced them to a pure language model (GPT-2-medium). We focused on GPT-2 L16 based on independent fMRI research (Caucheteux et al. 2022, see also **Supp Fig 5** for validation of this choice). Consistent with EEG reflecting traces of lexical processing we found that late linguistic Whisper layers captured all variance predicted by GPT-2 (and more) whereas earlier speech-like layers were complemented by GPT-2. **Mid-left:** To examine whether Whisper’s accurate EEG predictions were driven by contextualized representation, Whisper’s context window size was constrained to different durations [0.5s, 1s, 5s, 10s, 20s, 30s]. Accuracy was greatest at 10s, suggesting that intermediate contexts – which could support extraction of semantics and syntax – are valuable. Accuracy was on average ∼20% weaker for both shorter and longer contexts than 10s. Corresponding signed ranks test Z and FDR corrected p-values are displayed on the plot. **Mid-right**: To examine whether Whisper L6’s accurate EEG predictions were part driven by sub-lexical structure residual in Whisper’s last layer, we disrupted within word structure by either feature-wise averaging L6 vectors within word time boundaries or randomly reordering Whisper vectors within words. The outcome suggested that the EEG data additionally reflected a sub-lexical transformational stage, because either shuffling vectors within words or averaging them compromised EEG prediction in most participants. **Right**: To explore how the relative timing of EEG responses predicted by Whisper compared to the speech envelope and language model, we ran a battery of “single time lag” regression analyses. Model features were offset by a single lag within the range [0 to 750ms in 1/32s steps] and model-to-EEG mappings were separately fit on each lag, as repeated for each model in isolation. Whisper preferentially predicted a lag of 63ms, after the speech envelope (31ms) and before both the language model (125ms) and word surprisal (406ms). Note that the illustrated profiles chart (single-lag) prediction accuracies, and as such should not be confused with time-lagged regression beta-coefficients commonly used in the literature to estimate brain temporal response functions.

## Interpretation: EEG Preferentially Reflects the Encoding of 10s Speech Contexts

Because long multi-word speech contexts would be essential for Whisper to extract meaning and syntax from speech (and make associated predictions), we examined how valuable Whisper’s 30s context was for EEG prediction. To this end we generated Whisper L6 vectors using sliding context windows that we constrained to different durations [.5s 1s 5s 10s 20s 30s] to restrict the linguistic information Whisper might extract. As is illustrated in Figure 4 **Mid Left**, the strongest prediction accuracies were observed for 10s of context, and these accuracies were significantly greater than all other context window sizes except for 20s. However, although significant, the gain in prediction accuracy between 0.5s (Mean r=0.051) and 10s (Mean r=0.057) was modest, equating to 22% extra variance predicted (r^2^). Critically, this provides evidence that EEG signals are sensitive to multi-word speech contexts, which is essential for extracting and predicting linguistic representations and also something that is lacking in traditional context-invariant categorical speech models.

## Interpretation: EEG and Deep Whisper Layers Also Partially Reflect Sub-Lexical Structure

Because the bilateral temporal electrodes best captured by Whisper (Figure 2**/3**) are typically considered to reflect low-level speech acoustics and/or categorical speech units (DiLiberto et al. 2015 etc), and Whisper’s current EEG predictions could be partly driven by sub-lexical structure residual in Whisper (as was already hinted by the strong predictions obtained when pairing shallow speech-like whisper layers with GPT-2 in Figure 4 **left**), we tested this further. To this end, we evaluated whether Whisper L6’s predictive advantage was entirely lexically driven. If this was the case, we reasoned that either “lexicalizing” L6 by pointwise averaging vectors within word time boundaries, or randomly shuffling the temporal order of vectors within words should have negligible impact on EEG prediction accuracy.

To establish the effects of disrupting within word structure, we lexicalized or shuffled Whisper L6 vectors within words, as described above. We then ran comparative cross-validation analyses, first predicting EEG data with the Union of [Whisper L6 Env&Dv Lexical Surprisal] and then repeating analysis, but replacing Whisper L6 with either its shuffled or lexicalized counterpart.

Consistent with the EEG data also reflecting sub-lexical structure, both experimental manipulations of Whisper damaged prediction accuracy for most participants (Figure 4 **Mid Left**). Specifically, signed ranks comparisons of scalp-average prediction accuracies between Whisper L6 and lexicalized Whisper L6 revealed a significant drop in the latter (Mean±SEM=0.059±0.004 and 0.056±0.004 respectively, Z=2.86, p=0.0043, n=19, 2-tailed). When Whisper L6 vectors were randomly shuffled within words, and this process was repeated 20 times, the unshuffled prediction accuracies were found to be greatest in 13/19 participants. The cumulative binomial probability of achieving this outcome (p=1/20) in 13 or more participants is 2.5e-13. In sum, these analyses suggest that the current EEG correlates of speech-to-language transformation reflect a mixture of both lexical and sub-lexical structure.

## Interpretation: Whisper best predicts EEG responses that are intermediary between speech acoustics and language

Given the current evidence that EEG responses captured by Whisper reflect both lexical and sub-lexical structure we further examined how their timing related to responses to acoustic speech processing and language with the natural expectation that Whisper would be intermediary. To explore this, we ran a set of single time lag model-to-EEG prediction analyses in which model features were offset by only a single lag within the range [0 to 750ms in 1/32s steps] rather than the entire complement of lags as used in our other analyses. We reasoned that prediction accuracies derived from different lags would provide an estimate of the EEG response time-delay associated with each model. To simplify this analysis, model-to-EEG mappings were fit on isolated models (not model unions).

**Figure 4 (Right)** illustrates the cascade of EEG responses timings preferentially predicted by the different models (scalp-average prediction accuracies). Whisper preferentially predicted EEG at a time delay of 63ms which was indeed intermediary between the speech envelope (31ms) and the language model (GPT-2 L16 activation, 125ms) and word surprisal (406ms). This arrangement matches that sound->sub-lexical->lexical order that we naturally had expected. A secondary observation was that GPT-2’s L16’s prediction accuracy profile was doubled humped across response lags, with the second (weaker) peak at 563ms. We speculate the double hump reflects EEG responses associated with consecutive words e.g. the language model at word n both predicts the EEG response to word n and also n+1 (albeit with reduced accuracy).

As a side note, in interpreting Figure 4 **(Right)** please bear in mind that the single-lag prediction accuracy profiles are not equivalent to and should not be confused with the time-lagged regression beta-coefficients used frequently in the literature to illustrate neural temporal response functions (see Crosse et al. 2016 for examples). Please note also, that GPT-2 and probably Whisper (as above) are anticipatory, and may capture brain responses at short-latency.

## Interpretation: How Whisper Differs from Self-Supervised Speech Models that Infer Linguistic Representations

To gain further insight into the nature of EEG predictions based on Whisper’s explicit transformation of speech-to-language, we ran comparative analyses against two self-supervised speech models, that are trained entirely on sound data, with no access to language. For this, we selected Wav2Vec2 (Baevski et al. 2020) and HuBERT (Hsu et al. 2021) that in different studies have provided high accuracy fMRI models (Vaidya et al. 2022, Millet et al. 2022) and a strong basis for decoding model features from MEG and/or EEG data (Défossez et al. 2022, Han et al. 2023).

Specifically, like Whisper, Wav2Vec2 and HuBERT deploy Transformer encoders to build contextualized representations of speech, but different to Whisper they are pre-trained to infer the identity of artificially masked speech sounds. As such, the way these models represent speech and language should differ across their layers compared to Whisper. Specifically, past modelling and fMRI research has suggested that the inner layers of self-supervised speech models induce lexical semantic representations from speech (Pasad et al., 2021; Vaidya et al. 2022) – which could help them to infer contents of masked speech. However, the later layers focus on decoding back to an acoustic speech representation. As such, one might expect a different profile of EEG predictions across the layers of these models compared to the profile we have observed with Whisper – with any such differences adding to our understanding of the Whisper-based EEG predictions. To enable the current results to be cross-referenced to Millet et al. 2022 and Vaidya et al. 2022, we undertook EEG analyses with Wav2Vec2-base and HuBERT-base, which both have 12 layers, and compared them to Whisper-small (which has 12 rather than 6 layers). All networks were run with 30s sliding context widows, and all layer representations were projected onto ten corresponding PCA axes derived from an external dataset (7M, 7F speakers as before).

We found that all three speech models yielded highly accurate scalp-average EEG predictions (Figure 5). Whisper L12 was superficially the most accurate (Mean±SEM r=0.057±0.004), but when compared to the closest runner up Wav2Vec2 L7 (Mean±SEM r=0.055±0.004), there was no significant difference (Signed-rank Z=0.55, p=0.55, n=19, 2-tail). From visual inspection, the most salient difference between self-supervised models and Whisper was the pattern of EEG prediction accuracies across layers. As expected, whereas Whisper was characterized by an approximately monotonic increase in prediction accuracy tracking with layer depth, it was the intermediate layers of Wav2Vec2 and HuBERT (L7, L9 respectively) that were most accurate. Wav2Vec2 and HuBERT’s inner layer predictive advantage observed here on EEG closely mirrors the inner-layer advantages observed in fMRI research (Millet et al. 2022, Vaidya et al. 2022). Because comparative modelling analyses (Pasad et al. 2021) have specifically linked the representational content of Wav2Vec2 L7-8 to a lexical semantic model (GloVe Pennington et al. 2014), we presume this information supports EEG prediction. We further presume that this lexical content diminishes in latter model layers where it is back transformed to make predictions about masked sounds, and this accounts for the associated drop in EEG prediction accuracy.

**Figure 5.**
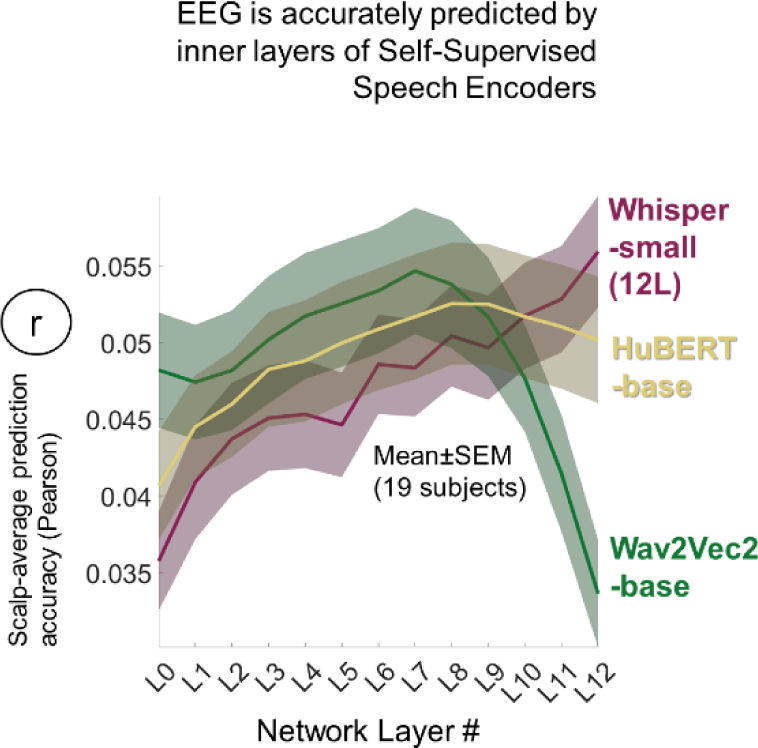
To explore how Whisper’s accurate EEG predictions compared to self-supervised speech models trained without direct access to language (on identifying masked speech sounds) we also repeated analyses with Wav2Vec2 and HuBERT. To enable cross-referencing to comparative fMRI studies (Millet et al. 2022, Vaidya et al. 2022), we performed this comparison on models comprising 12 layers. Wav2Vec2 and HuBERT both yielded highly accurate predictions but unlike Whisper, the inner layers were accurate. Further tests reported in the main text, suggest that Whisper predicted some different components of EEG signal to Wav2Vec2.

Finally, to explore for representational commonalities between Whisper L12 and Wav2Vec L7, we predicted EEG with the Union of the two models, and then contrasted accuracies with constituent models. Consistent with Whisper and Wav2Vec2 both contributing to prediction, the Union yielded modestly stronger prediction accuracies (Mean±SEM r=0.06±0.004, n=19) than both Wav2Vec2 L7 (Signed-Rank Z=3.67, p=2.5e-4, n=19, 2-tail) and Whisper L12 (Signed-rank Z=2.37, p=0.02, n=19, 2-tail). A variance partitioning analysis suggested that ∼75% of predicted variance was shared between models: r^2^_Shared_=100*(r^2^_Whisper_+r^2^_Wav2Vec2_–r^2^_Union_)/r^2^_Union_), ∼15% is added by Whisper L12: [100*(r^2^_Whisper_–r^2^_Shared_)/r^2^_Union_] and ∼9% is contributed by Wav2Vec2 L7: 100*(r^2^_Wav2Vec2_– r_2Shared_)/r_2Union_.

In sum, these findings reveal that EEG data is also accurately predicted by the more linguistic inner-layers of self-supervised speech models, and inner layers appear to share representational content with Whisper’s final most linguistic layer. Whisper’s unique contribution to EEG prediction may stem from its direct mapping to language in training. Collectively results are consistent with EEG reflecting a contextualized transformation of speech-to-language.

## Discussion

The current study has revealed electrophysiological correlates of the linguistic transformation of heard speech using a deep-learning approach – Whisper – that models speech-to-language transformation. This model addresses a key limitation of previous work that has typically invoked hand-crafted categorical speech units such as phonemes as an intermediary phase, and thereby neglected modelling the mechanism that maps sounds to categorical units. Given the current EEG results, this would appear to include neglecting to model the encoding of prior speech context. To strengthen the case that the newly predicted EEG signal reflects a linguistic transformation as opposed to a brain-like filter of concurrent acoustic speech, the study further demonstrated that the Whisper speech-to-language EEG response was sensitive to listener attention. Specifically, unlike EEG correlates of acoustic speech, the new EEG speech-to-language signature diminished when listeners ignored one speaker in favor of listening to another competing speaker (for whom the new EEG response was present). As a whole, this study exemplifies how deep-learning models can help tackle unresolved questions in human speech comprehension, and in so doing provide highly accurate predictions of neurophysiological data with minimal experimenter intervention.

The flipside of the above modelling benefits is that interpreting both the representations in deep-learning models and what they predict in brain data are both notoriously challenging. At the level of the deep-learning model, the inputs, outputs, architecture and training objective(s) provide clues to how information is reshaped throughout the network, and therefore what potentially could underpin EEG prediction. To recap Whisper is trained to translate multilingual speech spectra into words in either the input language or English and to also annotate word timing (see also **Methods**). The network architecture deploys a Transformer Encoder to contextualize and transform speech spectrograms into time-aligned vector representations (these vectors were the basis of the current study) and then a Transformer Decoder to transcribe the Encoder output vectors (NB we did not examine EEG using the Decoder to constrain the breadth of the current analyses). Thus, our starting assumption was that Whisper’s Encoder gradually transforms speech into a contextualized linguistic representation – which may be language invariant (perhaps encoding semantics) because the Encoder output is optimized to be decoded into words either within or between languages. However, at this stage the precise nature of network representations still remained ambiguous.

Having found that Whisper’s final “most linguistic” layer provided the most accurate EEG predictions across both electrodes and experiments, it was essential to run further interpretative tests to isolate what drove prediction (see also Vaidya et al. 2022). To probe for signatures of word processing we referenced Whisper’s EEG predictions to a pure language model (GPT-2, Figure 4 **Left**) - itself a Transformer Decoder - that predicts next-word identity. Consistent with EEG data mirroring elements of predictive lexical processing there were commonalities between Whisper and the language model, especially in Whisper’s final layer which captured all of the EEG variance predicted by the language model, and more besides. Interestingly, when the language model was paired with the earlier more speech-like Whisper layers, it did add complementary value, and generated EEG predictions on a par with latter Whisper layers. This suggested that EEG both reflects elements of word prediction (as found in GPT-2) alongside pre-lexical speech codes.

As a further probe of linguistic representation, we limited Whisper’s access to prior speech context, which should curtail Whisper’s ability to form semantic or syntactic representations and make predictions. Despite us having anticipated that EEG prediction accuracy might monotonically improve or asymptote with longer contexts (up to the 30s max), we found 10s context to be most accurate (∼20% extra variance predicted that .5s or 30s context). Critically the 10s contextual advantage provides evidence that EEG encodes multiword speech contexts. This gives reason to question the completeness of purely categorical speech models that are invariant to preceding lexical context. However, it remains unclear why a 30s context would be disadvantageous. Future work will be required to further characterize the neural correlates of contextualization and it may prove interesting to explicitly examine how context influences speech encoding at lexical and prelexical levels.

Relatedly we probed the EEG data for evidence of sub-word content (that could be residual in Whisper’s final layer) by disrupting within-word temporal structure (which should have little effect if entire words are represented with single codes). The findings were consistent with EEG additionally reflecting sub-word structure, as was evidenced by a modest reduction in EEG prediction accuracy in most participants. However, because EEG has low anatomical resolution, the degree to which this word/sub-word composite reflects intermediary part-speech part-language representational states in the brain, as opposed to a blurred sampling of distinct speech and language-selective neural populations is unclear from the current data. Related studies (Millet et al. 2022, Vaidya et al. 2022) referencing fMRI data (which has more precise anatomical resolution, but a slow sample rate) to self-supervised speech models such as Wav2Vec2 or HuBERT (which different to Whisper are trained on speech without access to language) or pure language models such as GPT-2 (Caucheteux et al. 2022) have provided evidence for graded cortical mappings with earlier model layers best predicting auditory cortices and deeper layers associated with language being more effective in regions radiating outwards across association cortices. Future work using intracranial recordings to explore superior temporal gyrus could be especially revealing, in light of past successes targeting fine-grained speech transformations (Mesgarani & Chang, 2014, Hamilton et al. 2021, Hamilton et al. 2018).

Finally, we ran comparative EEG analyses between Whisper – which explicitly transforms sound into language, and self-supervised speech models (Wav2Vec2 and HuBERT) – which have no access to language in training, but as it turns out induce linguistic representations in inner layers (Pasad et al. 2021, Vaidya et al. 2022) and therefore probably provide more reasonable models of early-stage language acquisition (Millet et al. 2022, Millet et al. 2021). Indeed, these inner language-like layers yielded the most accurate EEG predictions, which in the case of Wav2Vec2 were broadly equivalent to Whisper and overlapped in ∼75% of EEG variance predicted. The ∼15% extra variance predicted by Whisper may reflect Whisper’s direct transformation from speech to language. Nonetheless, these analyses provide convergent evidence that EEG reflects contextualized linguistic translation of heard speech.

To close, we believe the current study helps advance understanding of the neurophysiological processes underpinning speech comprehension and selective-attention. This adds to a growing body of research that has reaped the benefits of pre-trained deep-learning approaches for interpreting language or speech (Jain and Huth, 2018; Toneva and Wehbe, 2019, Sun et al. 2020, Anderson et al. 2021, Schrimpf et al. 2021, Millet et al. 2021, Heilbron et al. 2022, Millet et al. 2022, Vaidya et al. 2022, Antonello et al. 2023, Caucheteux et al. 2023, Goldstein et al. 2023). The current study has helped to broaden this horizon by identifying self-contained EEG model of speech-to-language transformation that is highly accurate, sensitive to listener attention and potentially revealing of how the brain could exploit prior speech context in comprehension. Because EEG which is both low-cost and widely available, and because the current speech-to-language model is automated, we hope that the approach can also contribute to applied research to help index the linguistic depth of speech processing in developmental and disordered populations.

## Methods

### EEG Data and Recording Parameters

All EEG data analyzed in this study are publicly available, and were downloaded from Dryad https://doi.org/10.5061/dryad.070jc. All EEG data were recorded from 128 scalp electrodes, plus two mastoid channels at 512Hz with a BioSemi ActiveTwo system. To emphasize the low frequency EEG signal that is commonly associated with prelexical and lexical representations (Di Liberto et al. 2015, Broderick et al. 2018), EEG was band-pass filtered between 0.5 and 8Hz using a 3^rd^ order Butterworth filter. Data were down-sampled to 32Hz to reduce the computational burden of forthcoming analyses. In all analyses, EEG data were re-referenced to the average of the mastoid channels.

### Single Audiobook EEG Experiment and Participants

Analyses presented in Figure 2 and **4** were undertaken upon EEG data originally recorded in (Di Liberto et al. 2015) from 19 participants (aged 19–38 years, 13 male) as they listened to an audiobook recording of The Old Man and the Sea (Hemingway, 1952) narrated by a male speaker. To mitigate participant fatigue, EEG recording was split into 20 runs, each of approximately 3mins duration, interleaved with brief breaks. The story line was preserved across the 20 trials, and the first trial corresponded to the start of the story.

### Cocktail Party Selective Attention EEG Experiment (Dual Audiobook) and Participants

Analyses presented in Figure 3 were undertaken upon EEG data originally recorded in Power et al. 2012 from 33 participants (aged 23–38 years; 27 male) who were played 30mins of two audiobooks presented simultaneously, but paid attention to only one book. The two audiobooks – 20,000 Leagues under the Sea (Verne, 1869) and Journey to the Center of the Earth (Verne, 1864) were narrated by different male speakers, and were presented via headphones, one to the left ear and the other the right. Participants were requested to selectively attend to one audiobook (in one ear) across the entire experiment. Audiobook presentation was split in thirty runs of 1 min each that were separated by brief breaks to mitigate participant fatigue. We analyzed 15 participants who paid attention to “20,000 Leagues…” and 12 who paid attention to “Journey…”. 6 Participants with incomplete EEG datasets were excluded to enable standardization of all analyses within the nested-cross validation procedure.

### Whisper – A Deep-Learning Model of Speech-to-Language Transformation

Our probe for EEG correlates of speech-to-language transformation – Whisper (Web-scale Supervised Pretraining for Speech Recognition or WSPSR/Whisper, Radford et al. 2022) – deploys a Transformer Encoder-Decoder deep-learning architecture that transforms spectral speech features into word transcriptions. Whisper is publicly available and we downloaded the version on Hugging Face (Wolf et al. 2019): https://huggingface.co/docs/transformers/model_doc/whisper) which was pre-trained to transcribe speech in 99 languages, to translate non-English speech to English words, and to annotate the timing of word boundaries.

Whisper’s Encoder and Decoder are both multilayer feedforward networks that each stack multiple layers of so-called Transformer blocks (e.g. L1-6 in Whisper-base, Figure 2**-4**, L1-12 in Whisper-small, Figure 5), where each block is itself a multilayer network. The Encoder receives a 30s speech spectrogram as input - an 80 channel Log-Mel Spectrogram, computed on 25ms windows, with step 10ms. The spectrogram is projected into a 512 dimensional space, operating on 20ms time-slices via a convolutional network (L0). Because Transformer layers process speech time-frames in parallel, a sinusoidal positional code is added to each time-frame to indicate its relative position in the sequence. The Transformer layers then refine each 512 dimension time-frame across successive layers into a time-aligned output vector sequence (also 512 dimensions). This output sequence is subsequently fed as a 30s whole into each Decoder layer (as a 1500*512 matrix, with each time-slice corresponding to 20ms.). The Decoder iteratively predicts the sequence of words corresponding to the Encoder output, one-by one, conditioned on the Encoder output, with each successive word prediction fed back as new Decoder input. Predicted words are represented as long vectors of probability values, with each entry corresponding to a single word. The current EEG analyses were exclusively based on Whisper’s Encoder module, and we excluded the Decoder to constrain the breadth of our analyses. We primarily focused analyses on Whisper-base rather than larger Whisper networks to reduce the computational burden of our analyses. Larger networks may produce more accurate EEG predictions (e.g. we observed predictions derived from Whisper-small to be more accurate Whisper-base, Figure 2 vs Figure 4).

The current analyses were based on three key assumptions: that Whisper’s Encoder transforms speech into representations that are (1) linguistic, (2) contextualized and (3) graded across network layers. Thus, successfully referencing EEG to Whisper would provide evidence that the brain signals reflect contextualized linguistic transformation of speech. Below we detail the bases of these assumptions.

Our assumption that Whisper’s Encoder generates linguistic representations was grounded on Whisper’s training objectives and architecture. Specifically, we assumed that the Encoder produces linguistic vectors, because they are shaped through optimization to be transcribed into words by the Decoder – optionally into a different language. Thus, Encoder vectors potentially suppress within and between speaker variation in intonation, accents, volume, intonation and so on to encode word identity in a language-invariant and possibly semantic format, at least to the extent these acoustic factors don’t compromise the Decoder. However, ultimately the precise linguistic nature of the Encoder vectors is an empirical question and one that we do pick up on in our latter Interpretative analyses with reference to a pure deep-learning language model (Figure 4 **Left**).

The assumption that Whisper produces contextualized speech representations is drawn from Whisper’s architecture, which preceded Whisper and was first introduced to generate contextualized language representations to support machine translation (Vaswani et al. 2017). For instance, without having some context to specify what the ambiguous English word “bat” refers to, the correct French translation would be unclear because French disambiguates the flying mammal (*chauvre souris* - bald mouse) from the sports tool (*batte*). In the case of modelling acoustic speech, linguistic context could be helpful to disambiguate noisy sounds, either within or across words (e.g. Diego Maradona played <NOISE>). Whisper’s Encoder’s Self Attention mechanism explicitly contextualizes each time-frame, by merging it with other time-frames via feature-wise weighted averaging (See **Supplementary Figure 1** for a detailed illustration, and **Figure 1**, and **Supplementary Figure 2 and 3** for a visualization of actual network weights). To provide some quick intuition, and taking an example from language, to encode “The vampire bat”, one would expect self-attention weights between “bat” and “vampire” to be strong (i.e. “bat” attends to “vampire” and the two become merged). Conversely attention weights between “bat” and “The” may be weak because “The” is not very informative for translating “bat”. Intuition aside, the value of contextualized speech encoding for EEG prediction is an empirical question which we address by experimentally constraining Whisper’s context (**Figure 4** **Mid Left**).

The third assumption - that Whisper produces a graded multistage transformation (of speech-to-language) - was based on Whisper’s multilayer feedforward architecture. Each layer (Transformer block) contextualizes each time-frame via the Self-Attention computation described above. Because in principle at least, L1-6 produce increasingly contextualized representations which contribute to transitioning speech into a linguistic code (suitable for Decoder transcription), we assumed that this transformation is graded. Nonetheless, we did experimentally test this assumption by referencing the EEG predictions derived from Whisper’s different layers to acoustic speech representations (**Figure 2**) and a language model (**Figure 4** **Left**).

For the EEG analyses we extracted 512 dimensional vectors corresponding to each (contextualized) time-frame, that were output from each layer. We focused only on encoder layer outputs rather than within-layer states (e.g. Self-Attention maps) to constrain the breadth of our analyses. As such, the current analyses may have overlooked representational information within Whisper that could be relevant for EEG prediction, and might provide the basis for future studies.

Because Whisper is set up to process 30s speech in parallel – which for the first time-frames in the current experimental stories would enable Whisper to see 30s into the future – we constrained the model to preserve biological realism. This was implemented by feeding the input speech audio waveform (resampled at 16000Hz) into Whisper via a ≤30s sliding window approach, which stepped forward over the waveform in 1/8s steps (corresponding to the 8Hz low-pass filtering of EEG data). Thus, at 10s into a run, Whisper was given access to the past 10s of speech audio waveform, and the series of Whisper vectors within the final 1/8s were saved from each layer of the Encoder and accumulated for the EEG analysis. At 40s into the analysis, Whisper processed 10-40s of audio waveform, and the series Whisper vectors within the final 1/8s were saved from each layer of the Encoder and accumulated for the EEG analysis. Whisper time-series were resampled from 50Hz to 32Hz (to match the EEG data) using the Python module resampy.

In addition, to reduce the computational burden of the EEG Regression Analysis, we applied Principal Components Analysis (PCA) to reduce the 512 dimensional Whisper vectors to 10 dimensions. To mitigate any risk of extracting data-set specific Principal Components and thereby maximize the generality of the approach, PCA axes were derived from an external audiobook dataset (i.e. not used as stimuli in any of the current EEG experiments). The external dataset collated 1.5mins speech from each of 7 males and 7 females narrating one story each (14 different stories in total). We processed each 1.5min dataset through Whisper separately, and extracted activations from L0-L6. We then concatenated each layer’s activations for the 14 stories to produce seven separate 512 *(14*1.5mins) matrices. PCA was conducted on each layer to provide 7 sets of PCA axes (L0 to L6). To then reduce the dimensionality of the Whisper datasets used in the current EEG analyses, Whisper vectors were projected onto the first ten Principal Component Axes of the corresponding layer by matrix multiplication. The selection of ten components was ultimately arbitrary, but was our first choice. To sanity check this choice, we ran a set of post hoc analyses with different numbers of components, and observed diminishing returns when using twenty or forty components (**Supplementary Figure 4**).

### Wav2Vec2 and HuBERT – Self-Supervised Models of Unlabeled Acoustic Speech

To explore how EEG correlates of Whisper relate to Self-Supervised Speech models that unlike Whisper are pre-trained entirely on unlabeled audio speech (without any access to language), we modeled audio stimuli with Wav2Vec2 (Baevski et al. 2020) and HuBERT (Hsu et al. 2021). Both Wav2Vec2 and HuBERT are publicly available, and we directly applied the base versions downloaded from https://huggingface.co/docs/transformers/model_doc/wav2vec2 or /hubert respectively.

Architecturally, both Wav2Vec2 and HuBERT share commonalities with Whisper in being contextualized speech models based on deep multilayer Transformer Encoder networks. However, unlike Whisper neither Wav2Vec2 or HuBERT uses a Transformer Decoder to transcribe Encoder outputs and indeed zero manually labeled data is used in pre-training. In the absence of having language to provide a ground truth for optimization, both Wav2Vec2 and HuBERT are pre-trained to make inferences about the identity of artificially masked speech sounds. For intuition, masking sound segments forces the Transformer blocks (and Self-Attention mechanism, see **Supplementary Figure 1**) to develop contextualized representations of the surround to infer the missing content. To facilitate inferring masked sound identity, both Wav2Vec2 and HuBERT are optimized to generate their own dictionaries of discrete speech sounds, which they approach in different ways (please see the original articles for further details of the differences in both quantization and training objectives). A final difference to Whisper is in audio pre-processing. Both Wav2Vec2 or HuBERT directly extract latent speech features from the audio speech waveform via a convolutional neural network, without spectrogram conversion.

Beyond the network differences, all other speech modelling parameters used for Whisper were carried over: Wav2Vec2 and HuBERT were run using the same ≤30s sliding window approach. Contextualized vector representations of each time-frame were extracted from each network layer (12 layers) and reduced to 10 dimensions via projection onto ten corresponding PCA axes, that had been separately computed for each layer on the same external audiobook dataset used in the Whisper analyses (14 stories narrated by 7 males and 7 females).

### GPT-2 – Modeling Word Prediction and Surprisal

To provide a pure language reference against which to interpret Whisper’s EEG predictions, we deployed GPT-2 (Generative Pretrained Transformer 2, Radford et al. 2019). GPT-2 is a Transformer Decoder Deep-Learning Model trained entirely upon language to predict the identity of the next word, given a history of up to 1024 preceding words. We selected GPT-2 based on its excellent performance in providing a basis for modelling brain data, spanning fMRI, MEG, ECoG and EEG (Goldsetin et al. 2022, Caucheteux et al. 2022, Heilbron et al. 2022, Schrimpf et al. 2021).

We deployed GPT-2 in two ways. The first was to capture the well-known N400 signature – a centroparietal negative response to unexpected words that is most pronounced at ∼400ms post word onset. Recent research has revealed that estimates of word surprisal (aka lexical surprisal) generated by GPT-2 provide an accurate way to recover the N400 response from continuous natural speech (Heilbron et al. 2022). The second application of GPT-2 was more exploratory in the context of EEG (though has been tested out in studies of fMRI, MEG and ECoG (Schrimpf et al. 2021, Caucheteux et al. 2022, Goldstein et al. 2022). Here we extracted word activation vectors output from each of GPT-2’s layers, and used these as a basis from predicting EEG activity in much the same way as we did for Whisper (please see the previous section).

Both word surprisal estimates and layer-wise word vectors were derived by processing the EEG stimulus story transcripts through GPT-2-medium (a 24 layer Transformer) using a sentence-based sliding window approach: GPT-2 was given as many previous sentences as could fit into its 1024 word context window, and the sliding window was advanced by stepping forward one sentence at a time. Beyond the transcript of the first EEG run, the sliding window straddled across run boundaries (so at the start of run 2, GPT-2 had access to sentences spanning backwards into run 1. We adopted this approach, because participants brains would likewise have had access to this prior story context. With each advance of the sliding window, word surprisal values and layer activations were extracted for all words within the leading (newest) sentence in the sliding window (detailed below).

Word surprisal estimates were computed for each word in the EEG stimulus, except for the very first word in the story (for which there was no preceding word from which to estimate surprisal). At word_n_, the estimate of next-word-identity is represented as a long vector of probability values linked to each word in GPT-2’s dictionary (this is GPT-2’s grand output). Word surprisal was computed as the negative log probability of the actual word_n+1_. To enable the EEG data to be referenced to the series of surprisal values, the surprisal values were aligned to word onset times as “spikes”. All other time-points were assigned the value zero.

To reference EEG to GPT-2 word activation vectors, we first harvested layer output vectors from each of the GPT-2’s 24 layers. Word vectors from all layers have 1024 dimensions. To reduce the computational burden of forthcoming Multiple Regression analyses, GPT-2 vectors were reduced from 1024 to ten dimensions through projection onto the first ten principal component axes derived from an independent storybook dataset (comprising the first 2250 words from 10 different stories). Ten dimensions were chosen to match the data reduction applied to Whisper. Exploratory analyses (not reported further) suggested there was no substantive advantage to using more than ten dimensions for predicting EEG data. The reduced 10 dimensional vector sequences were time-aligned to EEG, and vectors were stretched to fit to the duration of corresponding words (e.g. if word 3 started at 10s and ended at 10.3s, the vector for word 3 would be aligned to 10s and stretched to span 0.3s). Silent periods (lacking speech) were assigned the value zero.

As an addendum, to simplify the above explanation, we have implied that GPT-2 processes words. However more accurately, GPT-2 processes tokens which can either be words or sub-words (which can be useful to model out of dictionary words). For instance, the word “skiff” is treated as two tokens: “sk” and “iff”. In such a case GPT-2 would generate two token vectors for one word (or two surprisal estimates). In our analyses, the two token vectors would be combined into a single word vector by pointwise averaging – and the two token surprisal estimates would be averaged into a single word surprisal estimate.

### Speech Envelope and Derivative (Env&Dv) – A Model of Speech Audio Tracking

To model the EEG correlates of acoustic speech processing, we first computed the speech envelope, which is a time-varying measure of speech signal intensity, integrating across the acoustic frequency bands humans typically can hear. It is now widely accepted that cortical activity reflects the speech envelope (Aiken and Picton 2008; Destoky et al. 2019; Di Liberto, O’Sullivan, and Lalor 2015; Ding and Simon 2013; Etard and Reichenbach 2019; Lalor and Foxe 2010b; Nourski et al. 2009; Pasley et al. 2012). To compute the envelope, the speech audio waveform (44100Hz) was first lowpass filtered at 20 kHz (22050Hz cutoff frequency, 1 dB passband attenuation, 60 dB stopband attenuation). Then a gammachirp auditory filter-bank was deployed to emulate cochlea filtering (Irino and Patterson 2006) by filtering the 128 bands from 80 Hz to 8 kHz with an equal loudness contour. To create a unidimensional time-series, the 128 bands were averaged together.

Based on findings (Sohoglu and Davis 2020, Oganian and Chang 2019) that in addition to the speech envelope, cortical responses reflect so-called acoustic onsets – which correspond to positive slopes in the speech envelope – we computed this measure by differencing adjacent elements in the envelope time-series and reassigning negative values with zero.

For all EEG analyses we concatenated the Speech Envelope with acoustic onsets to produce a 2-dimensional time-series, abbreviated as Env&Dv (Dv because the latter measure reflects an approximation of the derivative).

### Log-Mel Spectrogram (Whisper’s Input)

As a further model of EEG correlates of acoustic speech processing, we deployed the 80 channel Log-Mel Spectrogram, that Whisper computes from the audio waveform preprocessing. The Log-Mel spectrogram is a re-representation of the Short-Time Fourier Transform Spectrogram emphasizing finer frequency resolution for lower frequencies and extracting signal amplitudes across a log-scaled filter bank (with more filters in low frequencies). Whisper’s implementation computes the Log-Mel Spectrogram using librosa: https://librosa.org/doc/main/generated/librosa.filters.mel.html, with 80 filters and an upper limit of 8000Hz.

### Mapping Models to Predict EEG with Multiple Regression and Nested Cross-Validation

To reference the continuous time-varying EEG signal back to either Whisper or the acoustic and lexical control models we used regularized multiple regression in a nested cross validation framework (**Figure 1**). Multiple regression was deployed to fit a predictive mapping from the time-aligned model/speech representation to each individual EEG electrode (repeated for all 128 electrodes).

To accommodate neural response delays, we temporally offset model/speech time-lines at each of a series of time-lags that stepped from the current (0ms) up to 750ms into the future (which would capture brain responses occurring up to 750ms post stimulation, which we assume spans the period in which speech is transformed to language). The profile of regression beta-coefficients over this 750ms period provides an estimate of the brains’ temporal response function (TRF) to stimulus features (See Crosse et al. 2016, for illustrations). To maintain consistency across all EEG analyses, the same time-lags were used for all models (whether acoustic/speech/linguistic) and combinations thereof.

Model-to-EEG TRF mappings were fit using Ridge Regression on EEG/model data for 18/20 runs. Both EEG and model/speech data sets (18/20 runs) were normalized by z-scoring as is commonplace for Ridge Regression, such that each feature or electrode had zero mean and unit standard deviation. Regression fitting was repeated for each of a range of different regularization penalties which can be considered to mitigate overfitting by squashing and smoothing potential outlier responses in TRF profiles to different degrees (lambda=[0.1 1 1e1 1e2 1e3 1e4 1e5]). The appropriate regularization penalty was estimated as the lambda value providing the most accurate EEG predictions of the 19^th^ “tuning” run, with accuracy averaged across all electrodes. To provide a final estimate of the TRFs ability to generalize to predict new data, the model-to-EEG TRF mapping corresponding to the selected regularization penalty was evaluated on the 20^th^ run. In tests on either the 19^th^ or 20^th^ run, prediction accuracy was evaluated separately for each electrode by computing Pearson correlation between the predicted time-series and the genuine EEG recording (see the circled r on **Figure 1**). Prior to this model/speech and EEG data for runs 19 and 20 were separately feature/electrode-wise normalized by z-scoring (see above). This procedure was repeated, whilst rotating the training/tuning/test run splits to generate separate prediction accuracy estimates for each of the 20 runs. To maintain consistency across all EEG analyses, the same Ridge Regression set up was deployed for all models (whether acoustic/speech/linguistic) and combinations thereof.

To summarize prediction accuracy across runs at each electrode, we computed the electrode-wise mean of correlation coefficients across the 20 runs (Correlation coefficients were r-to-z transformed prior to averaging then the mean was z-to-r back transformed by computing arctanh and tanh respectively). The scalp distribution of predicted-vs-genuine correlation coefficients provides a coarse estimate of which brain regions encode information found in the model/speech representation.

---

**Supplementary Figure 1 (Figure 1 Companion).**
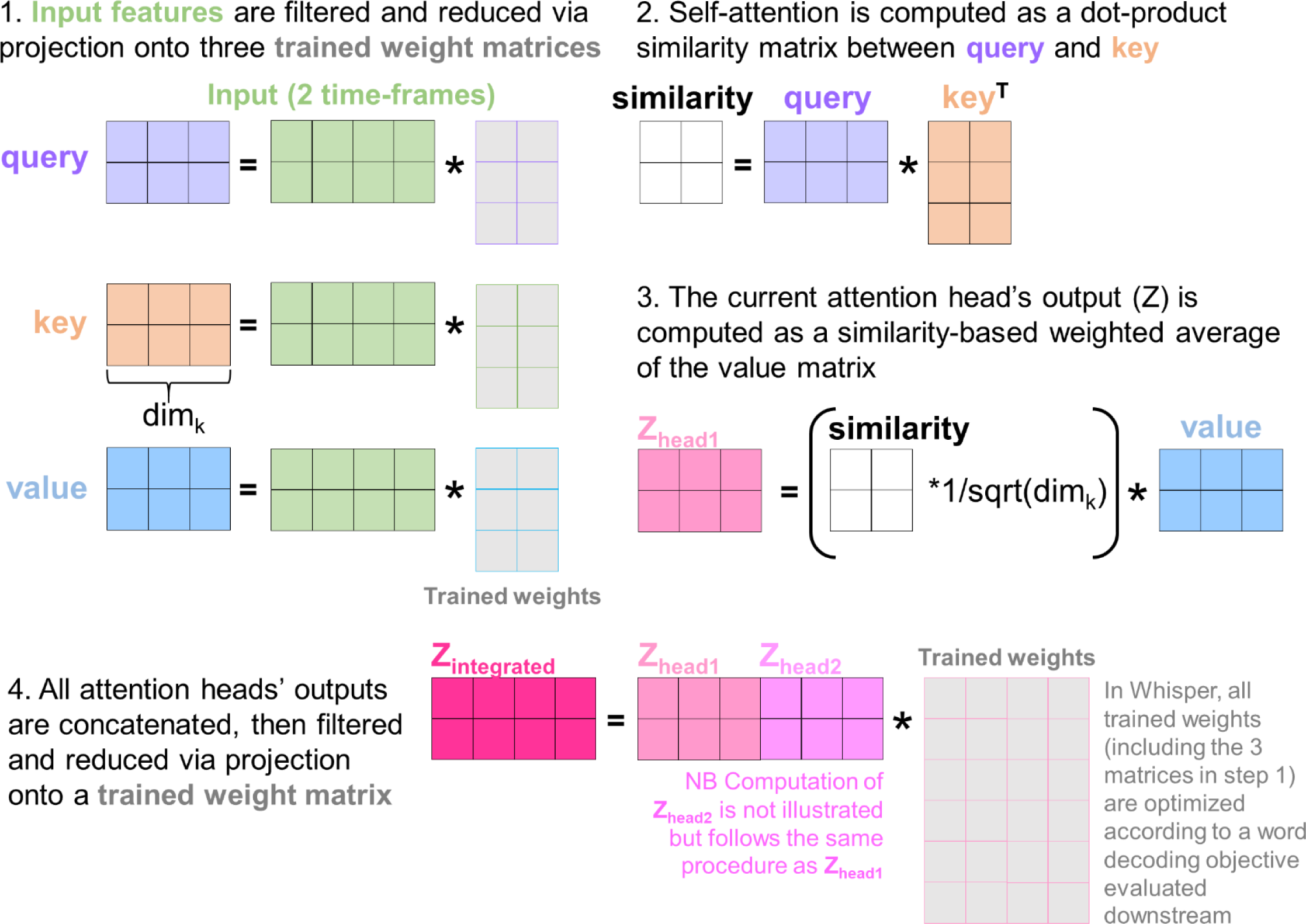
Illustration of the self-attention computation in a generic Transformer Encoder. The illustration borrows heavily from https://jalammar.github.io/illustrated-transformer/ and uses the same nomenclature and color-coding. To simplify visualization, computation in a single layer’s attention head is illustrated and layer inputs consist of only two time-frames containing four features. In practice the Whisper-base encoder has six layers, and eight separate attention heads per layer. Nonetheless, the computational procedure is the same for each layer and attention head. The initial transformer input (green) corresponds to the 80 channel Log-Mel Spectrogram, passed through a convolutional neural network, with positional codes then being added onto each time-frame to specify their relative order in the sequence. The input of successive layers is the output of the previous layer. At stage 1, information is extracted from the input, by projecting the input onto three separate trained weight matrices by matrix multiplication. The resulting representations are referred to as the query, key and value. The information extracted in the query and key matrices is critical for estimating the contextual relationships between different time-frames. At stage 2, contextual relations are estimated by multiplying the query with the transpose of the key. The resultant dot-product “similarity” matrix is populated by values that are high if query and key vectors resemble each other. The similarity matrix diagonal is liable to contain high values corresponding to the self-similarity between query and key vectors for the same time-frame. The attention head output Z is derived at stage 3. Z reflects the combination of the current time-frame with contextually-related time-frames – computed as a similarity-based weighted average of value vectors for all time frames. At stage 4, the outputs (Z) from all attention heads are concatenated, filtered and reduced through projection onto a trained weight matrix. The output Z_integrated_ is fed forward for subsequent processing before forming the layer output (see https://jalammar.github.io/illustrated-transformer/ for further details).

**Supplementary Figure 2 (Figure 1 Companion).**
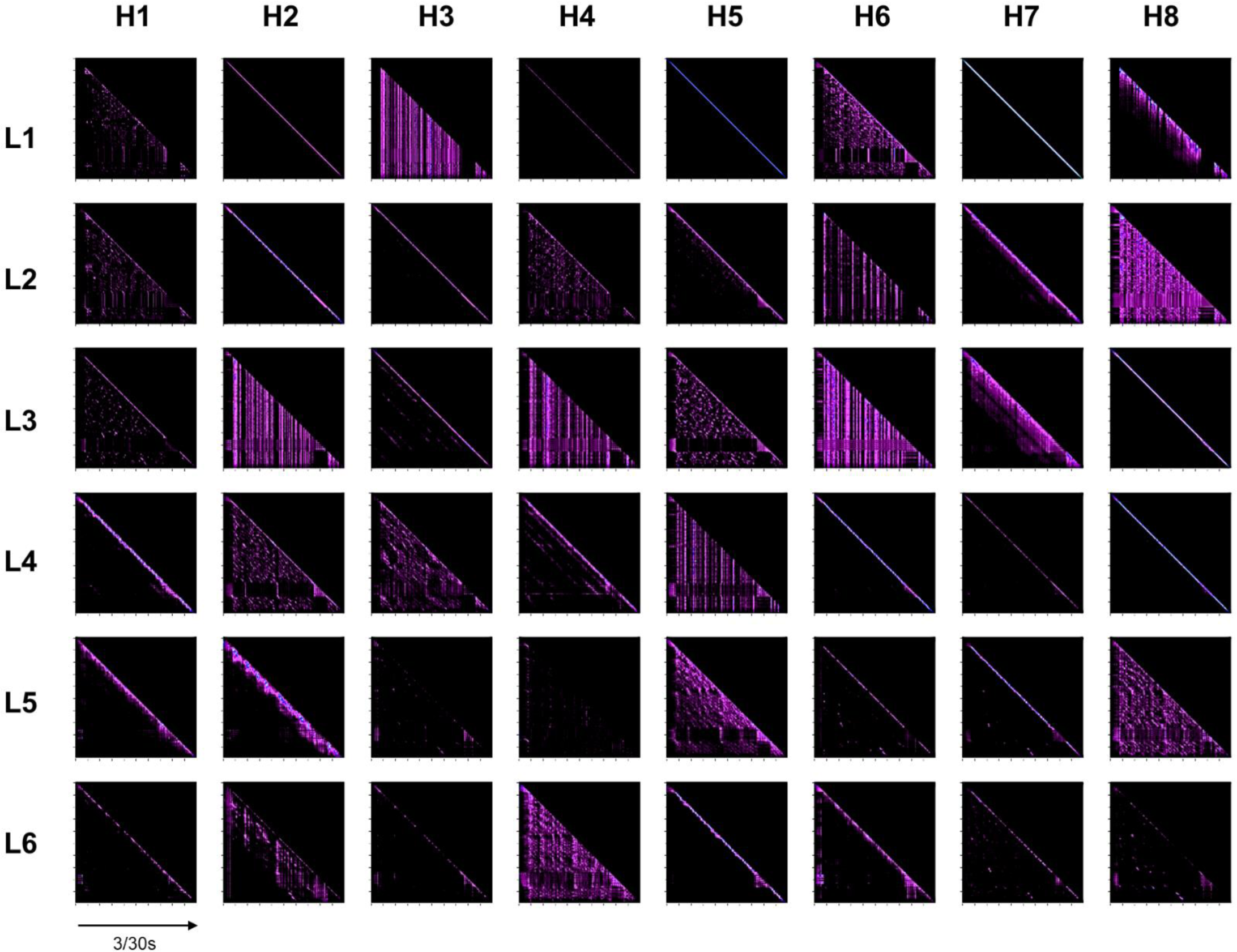
Comprehensive illustration of attention weights computed at each layer (L) and attention head (H), for the same 3s speech segment illustrated in Figure 1.

**Supplementary Figure 3 (Figure 1 Companion).**
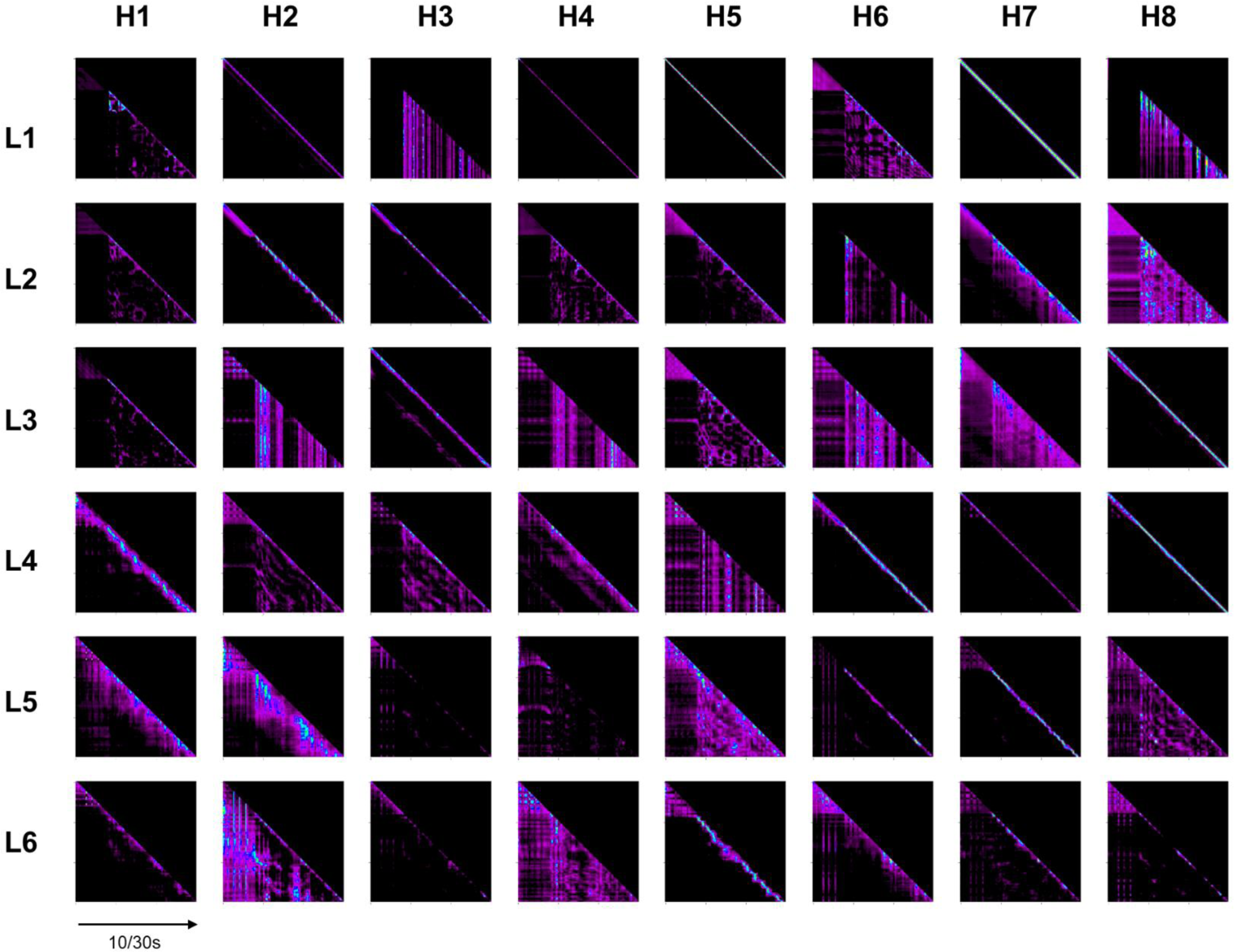
Comprehensive illustration of attention weights computed at each layer (L) and attention head (H), for a 10s speech segment, continuing on 7s after the 3s segment illustrated in in Figure 1, and Supplementary Figure 2.

**Supplementary Figure 4 (Figure 1 Companion).**
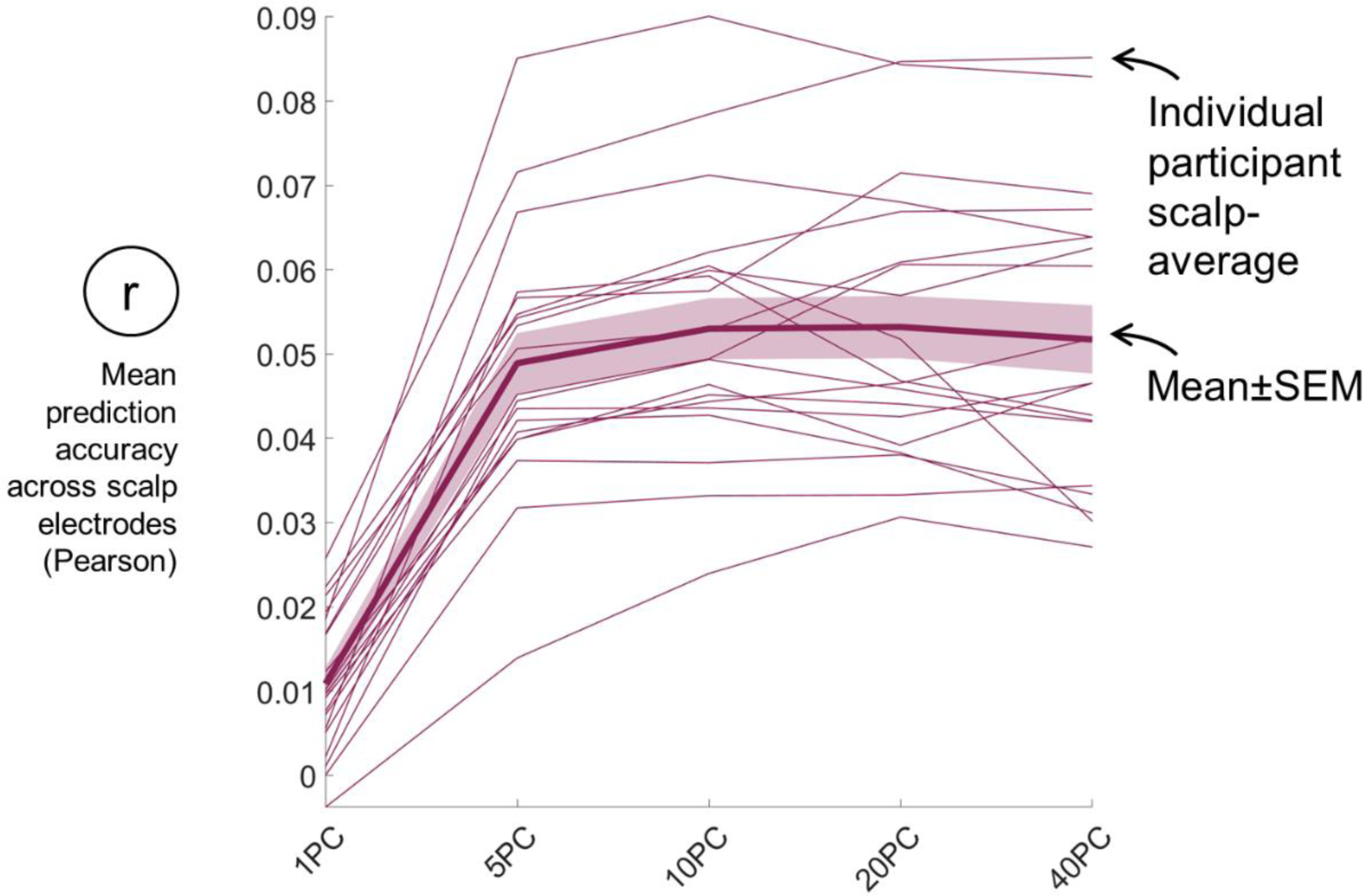
Exploration of how EEG prediction accuracy varies with the degree of data reduction applied to Whisper Layer 6. All Whisper-based analyses in the main article were performed following data reduction, as was achieved by projecting each Whisper layer onto a set of 10 principal component axes, that had been precomputed for each layer on Whisper representations derived from separate audiobook data. The selection of 10 components was our first choice, but arbitrary. To verify that the 10-component reduction was appropriate, the EEG data in Figure 2 was predicted after Whisper L6 had been reduced to [1 5 10 20 and 40] components. Visual inspection of scalp-average prediction accuracies (above) suggests that accurate EEG predictions could even be obtained with 5 components, and although the most accurate predictions were derived from 40PC, the performance boost above 10PC was not pronounced.

**Supplementary Figure 5 (Figure 4 Companion).**
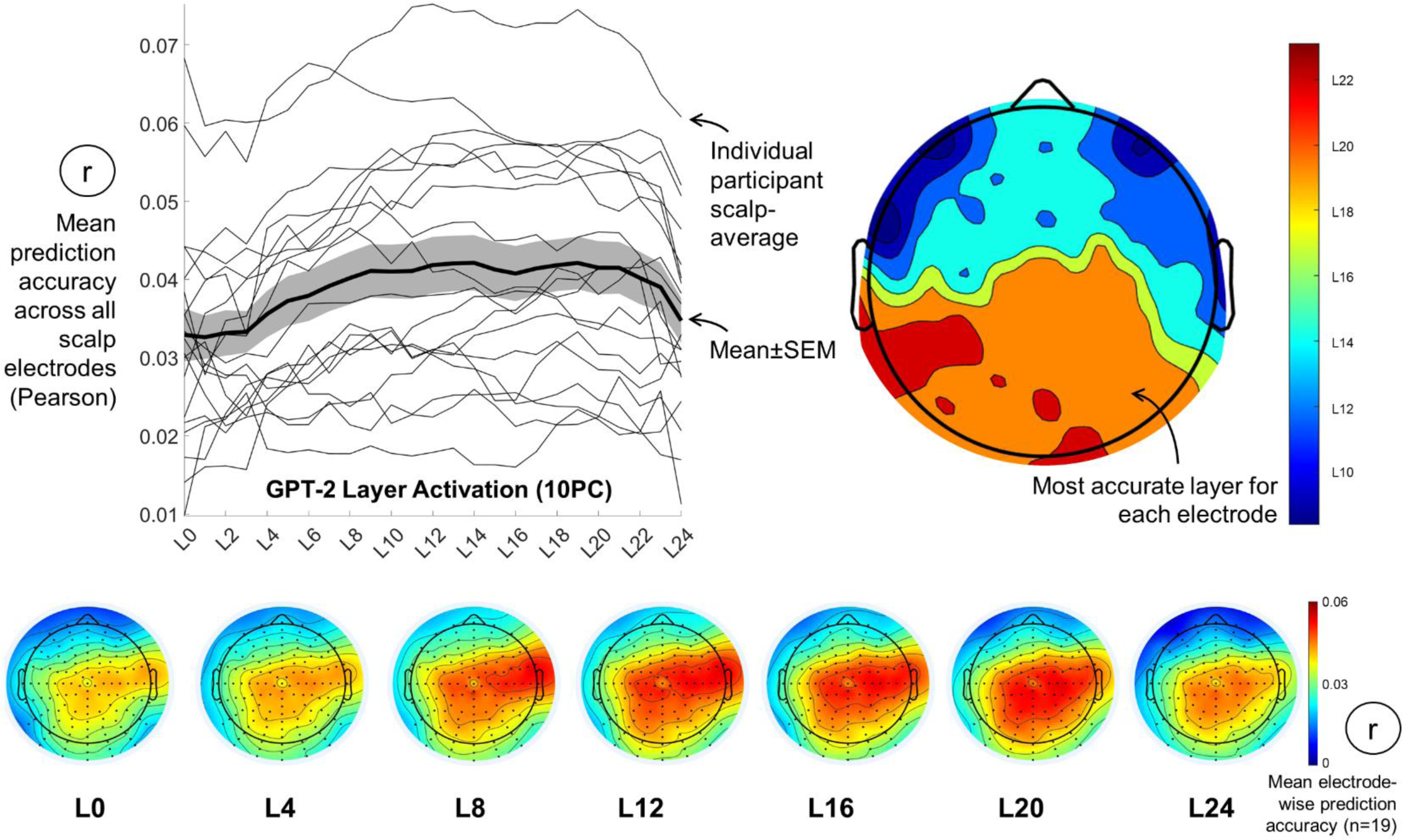
Exploration of how accurately representations at different layers of a language model (GPT-2-medium) predict natural speech EEG data Top left: Reanalysis of the audiobook EEG data from Figure 2 found that GPT-2 scalp-average EEG prediction accuracies were visibly greater for inner layers, mirroring independent analyses of natural speech fMRI data. Bottom: Electrode-wise prediction accuracies derived from successive GPT-2 layers. Top right: Scalp color-codes indicate the GPT-2 layer that most accurately predicted each layer. The most accurate layer was determined by (1) Computing the mean prediction accuracy across participants for each layer and electrode. (2) Identifying the layer yielding the maximum mean prediction accuracy at each electrode.

**Supplementary Table 1 (Figure 3 Companion).** Linear Mixed Effects analysis of the effects of Selective Attention on Layer-wise EEG Prediction Accuracies corresponding to the set of scalp-average prediction accuracies illustrated in **Figure 3** **Left Plots**. Model Fixed effects corresponded to Attention (nominal: Attended vs Unattended), the story heard (nominal: “Journey to…” or “20,000 Leagues…”), Whisper Layer (0 to 6), with interaction terms: Attention:Story and Attention:Layer and Attention:Story:Layer, and random effect of subject (nominal 1 to 27). The formula for the mixed model was: Accuracy ∼ 1 + Attention + Story + Layer + Attention:Story + Attention:Layer + Attention:Story:Layer + (1 | SubjectID). Outcomes are illustrated in the Table below. Most importantly the analysis revealed a significant interaction (p=4.5e5) between selective attention and Whisper layers (deep Whisper layers accurately predicted EEG, only if the modelled speech stream was attended). This interaction is visible in **Figure 3** as the positive trend between layer depth and prediction accuracy for attended speech, comparative to the weaker negative trend when speech is unattended. Otherwise, attention was the only significant main effect (p=1.84e-45: Prediction accuracies for attended speech were greater than unattended accuracies). Finally, the interaction between attention and story was significant (p=0.002). This was because for Journey to the Centre of the Earth, the boost in prediction accuracy from unattended to attended tended to be greater than corresponding values for 20,000 Leagues Under the Sea (Journey: Mean±SEM 0.023±0.004, n=12, 20,000: Mean±SEM 0.017±0.005, n=15, when scalp-average accuracies were averaged across all layers).

|  | Estimate | SEM | t | p |
| --- | --- | --- | --- | --- |
| **Intercept** | -0.39705 | 0.20002 | -1.985 | 0.047883 |
| **Attention (Nominal)** | 1.1747 | 0.072004 | 16.314 | 1.8346e-45 |
| **Story (Nominal)** | -0.19423 | 0.26836 | -0.72378 | 0.46966 |
| **Layer (0 to 6)** | -0.055444 | 0.050982 | -1.0875 | 0.27752 |
| **Attention : Story** | -0.29661 | 0.096604 | -3.0704 | 0.0022958 |
| **Attention : Layer** | 0.29762 | 0.0721 | 4.128 | 4.5263e-05 |
| **Story : Layer** | -0.036779 | 0.0684 | -0.5377 | 0.59111 |
| **Attention : Story : Layer** | 0.096022 | 0.096732 | 0.99266 | 0.32153 |

